# Development of a pan-neuronal genetic driver in *Aedes aegypti* mosquitoes

**DOI:** 10.1101/2020.08.22.262527

**Authors:** Zhilei Zhao, David Tian, Carolyn S. McBride

**Author notes:** Department of Integrative Biology and Museum of Vertebrate Zoology, University of California, Berkeley, CA 94720, USA.

## Abstract

The mosquito *Aedes aegypti* is the primary worldwide vector of arboviruses that infect humans, including dengue, Zika, chikungunya, and yellow fever. Recent advances in transgenic technology have yielded important new insight into the biology of this disease vector. The early development of neurogenetic tools, in particular, is beginning to shed light on the neural basis of behaviors that allow *Ae. aegypti* to thrive in human environments and find and bite human hosts. Despite these advances, a pan-neuronal expression driver remains elusive. Pan-neuronal drivers give researchers genetic access to all neurons and thus provide a critical entry point for circuit dissection. Here, we describe our efforts to generate pan-neuronal drivers in *Ae. aegypti* via targeted knock-in of in-frame reporter constructs to the native coding sequence of broadly expressed neural genes with CRISPR/Cas9. Two of five attempts were successful, resulting in a *Syt1:GCaMP6s* strain that expresses synaptically-localized GCaMP in all neurons and a *brp-T2A-QF2w* driver strain that can be used to drive and amplify expression of any effector in all neurons via the Q binary system. We show that both manipulations broadly and uniformly label the nervous system and have only mild effects on behavior. We envision that these strains will facilitate neurobiological research in *Ae. aegypti* mosquitoes and provide documentation of both successful and failed manipulations as a roadmap for similar tool development in other non-model species.

## Introduction

Mosquito-borne diseases are a major threat to public health, causing nearly a million deaths and hundreds of millions of non-lethal infections each year (WHO, 2020). One particularly dangerous mosquito is *Aedes aegypti*, the primary vector of dengue, Zika, chikungunya and yellow fever (Christophers, 1960). *Ae. aegypti* originated in Africa but spread rapidly across the global tropics and subtropics within the last 500 years, putting billions of people at risk (Powell et al., 2018; WHO, 2020). The success of this species is largely attributable to a rich repertoire of behaviors that adapt the mosquito to human hosts and habitats. Females experience a carefully regulated 3-4 day cycle during which they alternately seek humans for biting, resting sites for digestion, and containers of water for egg laying (Christophers, 1960). At each stage they must integrate and respond appropriately to a distinct set of sensory cues including chemical, visual, and thermal stimuli (Clements, 1999). Understanding the neural basis of these and other mosquito behaviors is both interesting from a neurobiological perspective and important for the design of effective and specific mosquito control strategies.

The recent optimization of CRISPR/Cas9 and other transgenic technology in *Ae. aegypti* (Kistler et al., 2015; Li et al., 2017; Nimmo et al., 2006; Häcker et al., 2017; Anderson et al., 2010; Kokoza et al., 2001) has opened the door to the development of powerful neurogenetic tools that promise to take our understanding of its behavior to the neural level. These tools include binary expression systems with cell-type-specific drivers (Kokoza and Raikhel, 2011; Matthews et al., 2019; Riabinina et al., 2016). Pan-neuronal expression drivers, however, remain elusive. Pan-neuronal drivers allow researchers to express a reporter or effector in all neurons and thus provide a critical entry point for circuit dissection. They are typically generated by fusing the promoter region of a broadly expressed neural gene to a reporter and inserting this ‘promoter fusion’ into a random location in the genome via a transposase or site-specific integrase (Figure 1A, left). The reporter may be the desired effector molecule, such as a fluorescent protein or calcium indicator, but it is more commonly the transcriptional activator from a binary expression system (Venken et al., 2011), designed to drive and amplify expression of an effector molecule located elsewhere in the genome.

**Figure 1.**
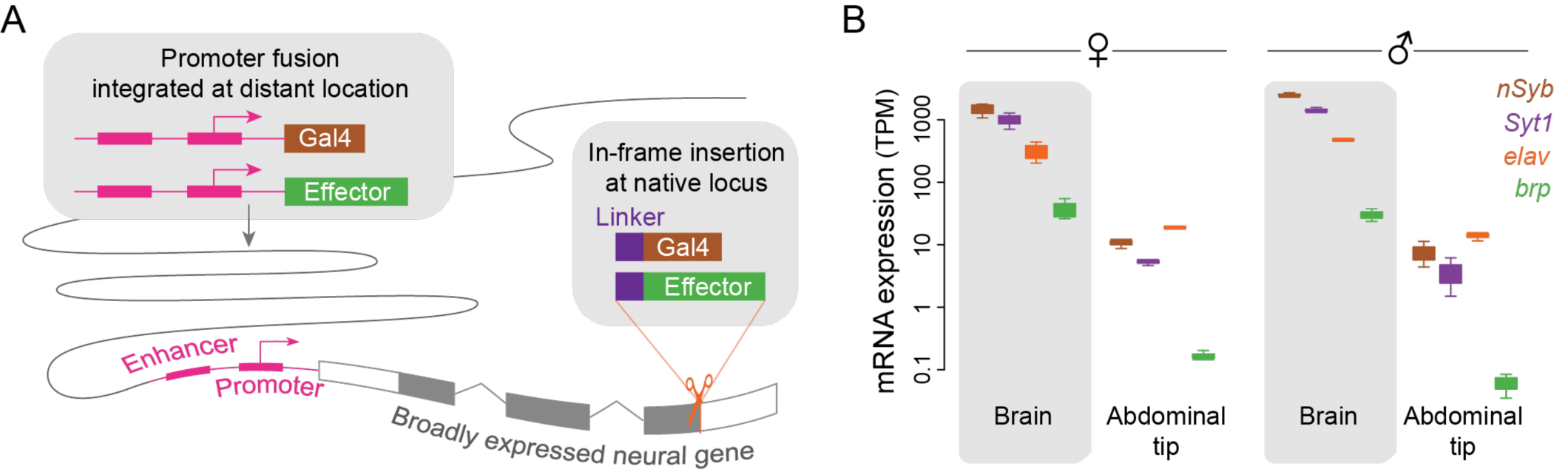
Two approaches to the generation of pan-neuronal drivers leverage the cis-regulatory elements of a broadly expressed neural gene. (**A**) In a promoter fusion (left box), several kilobases of sequence in the promoter region of the target gene is fused to an effector or transcriptional activator and inserted at a distant (often random) location in the genome. Alternatively (right box), the effector or activator is inserted in-frame into the target locus preceded by a short linker that can be engineered to result in translation of separate target and reporter proteins from the same transcript (e.g. T2A ‘ribosomal skipping’ sequence) or a fused protein (e.g. 3XGS linker). (**B**) Expression of candidate neural target genes in the brain and a non-neural tissue of *Ae. aegypti* (data from Matthews et al., 2016; n=3-8 RNAseq libraries per tissue). *nSyb, n-Synaptobrevin; Syt1, Synaptotagmin1; elav, embryonic lethal abnormal vision; brp, bruchpilot*.

Promoter fusions are widely used as expression drivers in the model insect *Drosophila melanogaster*. While there are also some successful examples in mosquitoes (Bui et al., 2019; Kokoza and Raikhel, 2011; Li et al., 2017; Papathanos et al., 2009; Riabinina et al., 2016), performance is generally inconsistent. In a recent study of the African malaria mosquito *Anopheles gambiae*, the promoter fusion for only one of four target genes was functional (Riabinina et al., 2016). Attempts to make pan-neuronal promoter fusions in *Ae. aegypti* have also been unsuccessful (Ben Matthews and Meg Younger *pers. comm*.), possibly because regulatory elements are often scattered across large intergenic regions in this species (Matthews et al., 2018; Nene et al., 2007). Recently, the *Ae. aegypti* polyubiquitin promoter was used to generate a promoter fusion line that expresses the calcium indicator GCaMP6s in all cells of the mosquito (Bui et al., 2019). This strain has been used successfully for calcium imaging (e.g. Lahondère et al., 2020; Melo et al., 2020), but the presence of GCaMP6s in all cells, not only in neurons, can complicate interpretation.

An alternative approach to the generation of genetic drivers that replicate the pattern of expression of a target gene is to insert an in-frame reporter construct directly into the target locus via homology directed repair (Figure 1A, right). Inclusion of a ‘ribosomal skipping’ sequence such as T2A before the reporter can be used to generate separate target and reporter proteins from the same transcript (Diao and White, 2012), while inclusion of a flexible linker (e.g. 3XGS) can be used to generate a fused protein that is localized according to signal sequences in the target protein. In-frame insertions take advantage of the cis-regulatory elements of the target gene *in situ*, avoid the positional effects inherent to random insertions, and were recently shown to be effective in *Ae. aegypti* (Matthews et al., 2019). While it is important to keep in mind that these manipulations alter the target locus, harmful effects may be minimized by placing the insertion at the very end of the coding sequence (to preserve the target protein) and/or by carrying out experiments in heterozygotes.

Here, we use targeted insertions to generate two pan-neuronal strains in *Ae. aegypti* mosquitoes. The first strain expresses synaptically localized GCaMP in all neurons for use in neural imaging (*Syt1:GCaMP6s*). The second is a flexible pan-neuronal driver that can be used in concert with the Q binary expression system to drive the expression of any effector in neurons (*brp-T2A-QF2w*). We also describe several failed attempts made during our troubleshooting process, which we hope will be informative for similar efforts in this and other non-model organisms.

## Results

### Identification of neural genes for targeted knock-ins

We first set out to identify a set of broadly expressed neural genes to target with knock-in constructs. Previous work in *Drosophila* pointed to four candidates: *neuronal Synaptobrevin* (*nSyb*), *Synaptotagmin1* (*Syt1*), *bruchpilot* (*brp*) and *embryonic lethal abnormal vision* (*elav*). The first three genes encode proteins involved in the maintenance and structure of chemical synapses (Südhof, 2012), while *elav* encodes a protein involved in neuron-specific mRNA splicing (Yao et al., 1993). In *Drosophila*, all four are thought to be expressed in almost all neurons (but see Davis et al., 2020; Takemura et al., 2008) and not in other cell types. The promoter regions of *nSyb* and *elav* were used to generate the most popular pan-neuronal drivers available in *Drosophila* (Luo et al., 1994; Pauli et al., 2008; Riabinina et al., 2015). We identified the *nSyb, Syt1, brp*, and *elav* orthologs in the *Ae. aegypti* reference genome (Matthews et al., 2018) by BLAST homology to the *D. melanogaster* proteins and queried previously published RNAseq data to confirm that expression was highly enriched in the *Ae. aegypti* brain compared to a mostly non-neural tissue (abdominal tip) (Figure 1B). While all four genes are presumably co-expressed in the same cells, they varied significantly in absolute expression. Notably, *nSyb* and *Syt1* transcripts were 30-60 times more abundant than *brp* transcripts (Figure 1B). We designed and tested single guide RNAs (sgRNAs, n=3-6 per gene) to direct *Cas9* nuclease to sequences near the stop codon of each gene. We identified sgRNAs that cut efficiently for both *Syt1* and *brp*, but not for *nSyb* and *elav*.

### Generation of a pan-neuronal, synaptically localized GCaMP line

Our initial goal was to generate a mosquito line with pan-neuronal expression of the calcium indicator GCaMP for neural imaging applications. We decided to start with a direct, in-frame insertion of GCaMP6s (Chen et al., 2013) into the *Syt1* locus. While simple, this strategy directly ties the level of GCaMP6s expression to that of the target gene, without the amplification inherent in binary expression systems. To ensure adequate GCaMP6s expression, we therefore chose to target *Syt1* due to its high expression (Figure 1B), and to insert three tandem copies of GCaMP6s instead of one (*Syt1-T2A-3XGCaMP6s*, Figure 2A). The GCaMP6s effectors were separated from each other and from the *Syt1* coding sequence by T2A motifs in order to generate up to four separate proteins (1 Syt and 3 GCaMP6s) from the same transcript. We targeted the last exon of *Syt1*, seven codons upstream from the stop codon, and preserved the native protein by including those final codons in the insertion. This final exon is shared among all splice forms according to the most recent annotation of the *Ae. aegypti* genome (NCBI LOC5565901).

**Figure 2.**
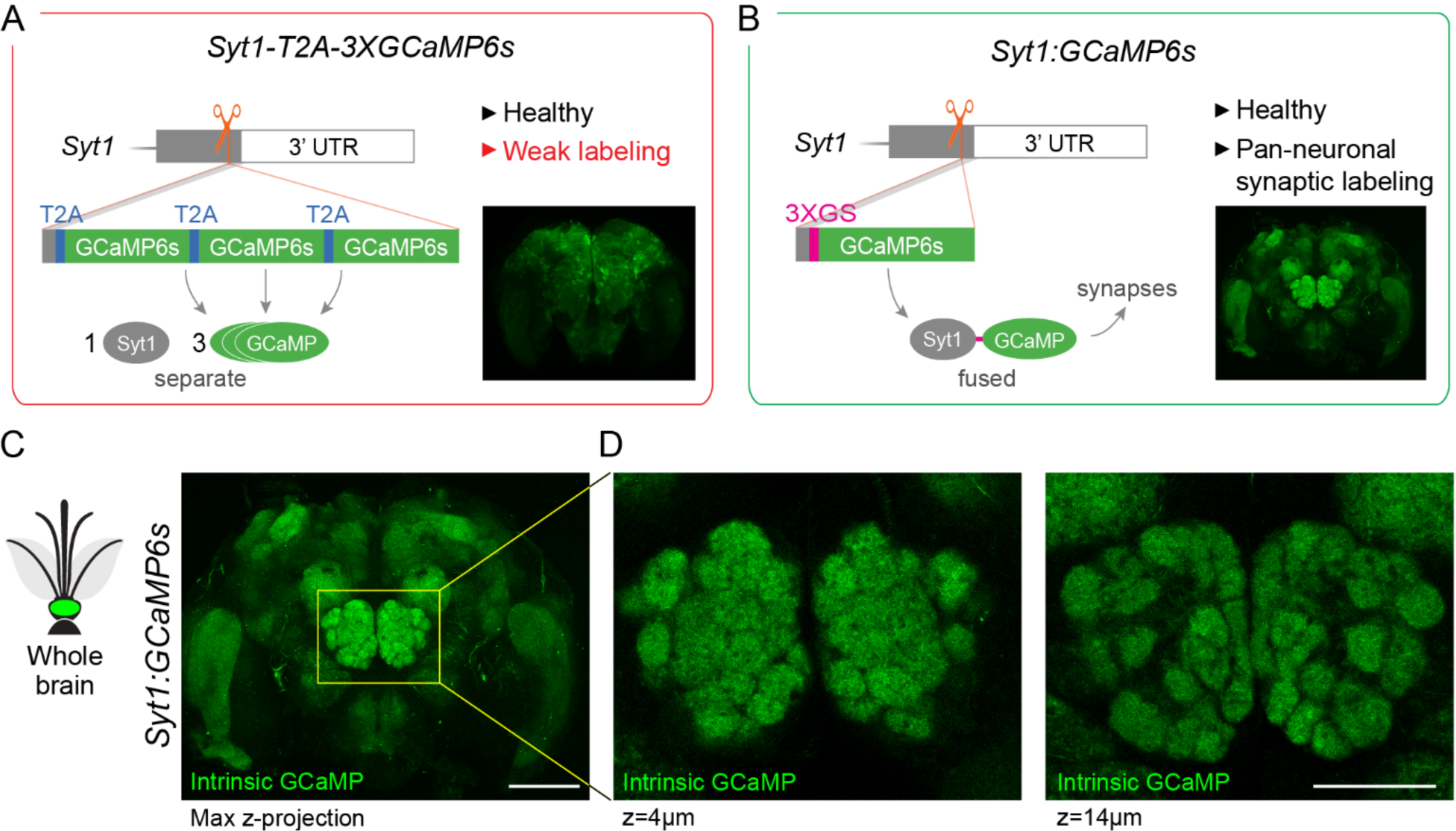
Generation of a pan-neuronal, synaptically localized GCaMP line for neural imaging in *Ae. aegypti*. (**A**,**B**) Summary schematics for two alternative strategies showing construct design and outcome (red border indicates failure for reason given in red lettering; green border indicates success). The first strategy (**A**) involved in-frame insertion of three copies of GCaMP6s separated by T2A ribosomal skipping sequences designed to generate separate Syt1 and GCaMP proteins (*Syt1-T2A-3XGCaMP6s*). Expression appeared pan-neuronal (inset), but GCaMP expression was too weak for neural imaging. The second strategy (**B**) involved in-frame insertion of GCaMP6s preceded by a 3XGS linker designed to generate a Syt1-GCaMP fusion protein (*Syt1:GCaMP6s*). By concentrating GCaMP at presynaptic sites, this approach produced a healthy line bright enough for imaging. Insets show anti-GFP staining (**A**), or intrinsic GCaMP fluorescence (**B**) in adult brains. Both constructs also included a screening marker (3XP3-dsRed, not shown). (**C**,**D**) Intrinsic GCaMP fluorescence in brain (**C**) and antennal lobe (**D**, 2 z-planes) of *Syt1:GCaMP6s* heterozygous male. Scale bars 100 µm (**C**), 50 µm (**D**).

The *Syt1-T2A-3XGCaMP6s* knock-in was successful in that we were able to isolate a stable line that showed broad expression of GCaMP6s in neurons of the mosquito brain (Figure 2A, inset). However, GCaMP6s expression was too weak for neural imaging (data not shown). In these mosquitoes, GCaMP6s molecules should be distributed throughout the cytosol. Endogenous Syt1 proteins, in contrast, are translocated to presynaptic sites. We therefore reasoned that by fusing GCaMP6s to Syt1 – rather than simply tying its mRNA expression to that of Syt1 – we could concentrate the limited supply of GCaMP6s at synapses and enhance brightness. Synaptically localized GCaMP6s also offers the ability to record activity selectively from presynaptic neurons (e.g. Cohn et al., 2015).

To test the fusion strategy, we again knocked GCaMP6s into the *Syt1* locus, this time replacing the T2A-3XGCaMP6s donor payload with a 3XGS flexible linker followed by a single copy of GCaMP6s (*Syt1:GCaMP6s*, Figure 2B). As expected, this line broadly and strongly labeled synapses throughout the brain (Figure 2C). Confocal imaging revealed strong intrinsic GCaMP6s fluorescence in all major neuropils, including the antennal lobe (Figure 2D). The antennal lobe is the primary olfactory processing center of insect brains and consists of spherical units of neuropil called glomeruli, where primary sensory neurons synapse onto second order projection neurons and interneurons (Vosshall and Stocker, 2007). The architecture of this area makes it an ideal location to check labeling since missing or unevenly labelled glomeruli are easily detected. We found that all glomeruli in the *Syt1:GCaMP6s* line were labeled with similar strength and that glomerular boundaries were clearly visible (Figure 2D). Preliminary 2-photon imaging experiments under reasonable laser power (<15mW) also revealed measurable and glomerulus-specific changes in fluorescence in response to single odorant stimuli (data not shown).

### Generation of a flexible pan-neuronal driver line

We next turned to the construction of a flexible pan-neuronal driver line that could drive expression of diverse effector transgenes in all neurons via a binary system. We decided to use the Q binary system, which was recently validated in both *Anopheles gambiae* and *Ae. aegypti* mosquitoes (Matthews et al., 2019; Riabinina et al., 2016) and again target the *Syt1* stop codon. More specifically, we generated an in-frame T2A-QF2 insertion to enable independent translation of the QF2 transcriptional activator in all neurons. Typically, one would then cross this driver to a second strain carrying a QUAS effector transgene. However, in order to quickly test the system in a single step, we included a QUAS-GCaMP6s effector in the knock-in construct, just downstream of QF2 (*Syt1-T2A-QF2-QUAS-GCaMP6s*, Figure 3A). This decision proved fortuitous. We were able to confirm strong, pan-neuronal expression of GCaMP in two families of G1 larvae (n>20 larvae total) while screening for transformants under an epifluorescence microscope (Figure 3A, inset), despite the fact that all labeled larvae arrested development at the 2nd or 3rd instar and eventually died.

**Figure 3.**
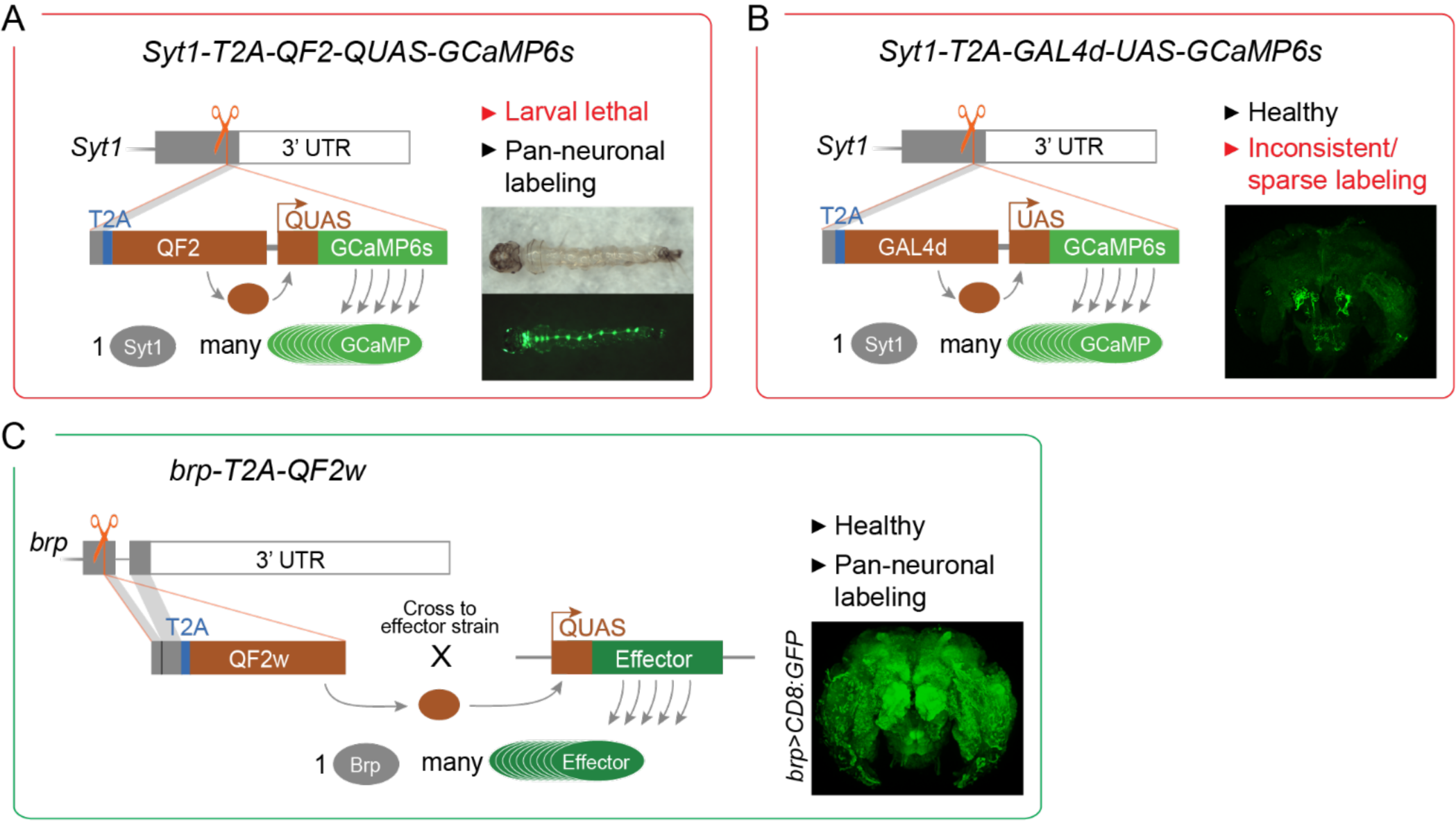
Generation of a pan-neuronal driver line in *Ae. aegypti*. Each panel provides a summary schematic for one of the three alternative strategies (red border, failed; green border; successful). (**A**,**B**) The first two strategies involved in-frame insertion of transcriptional activators near the end of the native *Syt1* locus designed to generate separate Syt1 and QF2 (**A**) or Gal4d (**B**) proteins. Corresponding GCaMP6s effector elements were included in tandem to enable rapid one-step testing of pan-neuronal expression. Insets show intrinsic GCaMP fluorescence in larva (**A**), or anti-GFP staining in adult brains (**B**). Both approaches failed for reasons provided (red font). (**C**) The third strategy involved in-frame insertion of the QF2w transcriptional activator near the end of the native *brp* locus, designed to result in more modest levels of expression of the weaker transcriptional activator. Inset shows anti-GFP staining in brain of adult female from cross between *brp-T2A-QF2w* and *QUAS-CD8:GFP*. All donor constructs also included a screening marker (3XP3-dsRed, not shown).

Both QF2 and GCaMP have been shown to cause fitness defects when expressed pan-neuronally (Riabinina et al., 2015; Steinmetz et al., 2017), and we suspected that overexpression of one or both was responsible for the observed larval lethality. We therefore decided to replace QF2 with Gal4d (and QUAS with UAS) to see whether a different binary system would solve the problem (*Syt1-T2A-GAL4d-UAS-GCaMP6s*, Figure 3B). We chose GAL4d instead of full-length GAL4 because it is substantially shorter (making the knock-in more efficient) and less potent in driving expression (reducing potential toxicity caused by GCaMP expression) (Ma and Ptashne, 1987; Pfeiffer et al., 2010). Unfortunately, however, while we successfully isolated healthy transformants, GCaMP labelling was sparse and highly variable across individuals (Figure 3B inset).

Full-length Gal4 has been shown to drive reporter expression in *Ae. aegypti* (Kokoza and Raikhel, 2011). It is therefore possible that the failure of the second driver strain resulted from our use of Gal4d (though others have also had problems getting the Gal4/UAS system to work in *Ae. aegypti*; Matthews et al, 2019). Nevertheless, we decided to return to the Q system and address the lethality problem in other ways (Figure 3C). To mitigate potential QF2 toxicity, we replaced QF2 with QF2w – a weaker and less toxic version of the same transcriptional activator (Riabinina et al., 2015). We also targeted *brp* instead of *Syt1*. Since *brp* is expressed 30-40 times less highly than *Syt1*, we expected this change to result in a substantial reduction in QF2w expression. Finally, to guard against potential toxicity of GCaMP6s, we removed the QUAS effector. The resulting knock-in strain (*brp-T2A-QF2w*, Figure 3C) was stable and produced viable, seemingly healthy adults both in isolation and when driving expression of *CD8:GFP, Syt1:tdTomato*, or *GCaMP6s* (Figure 4 and Jové et al., 2020).

**Figure 4.**
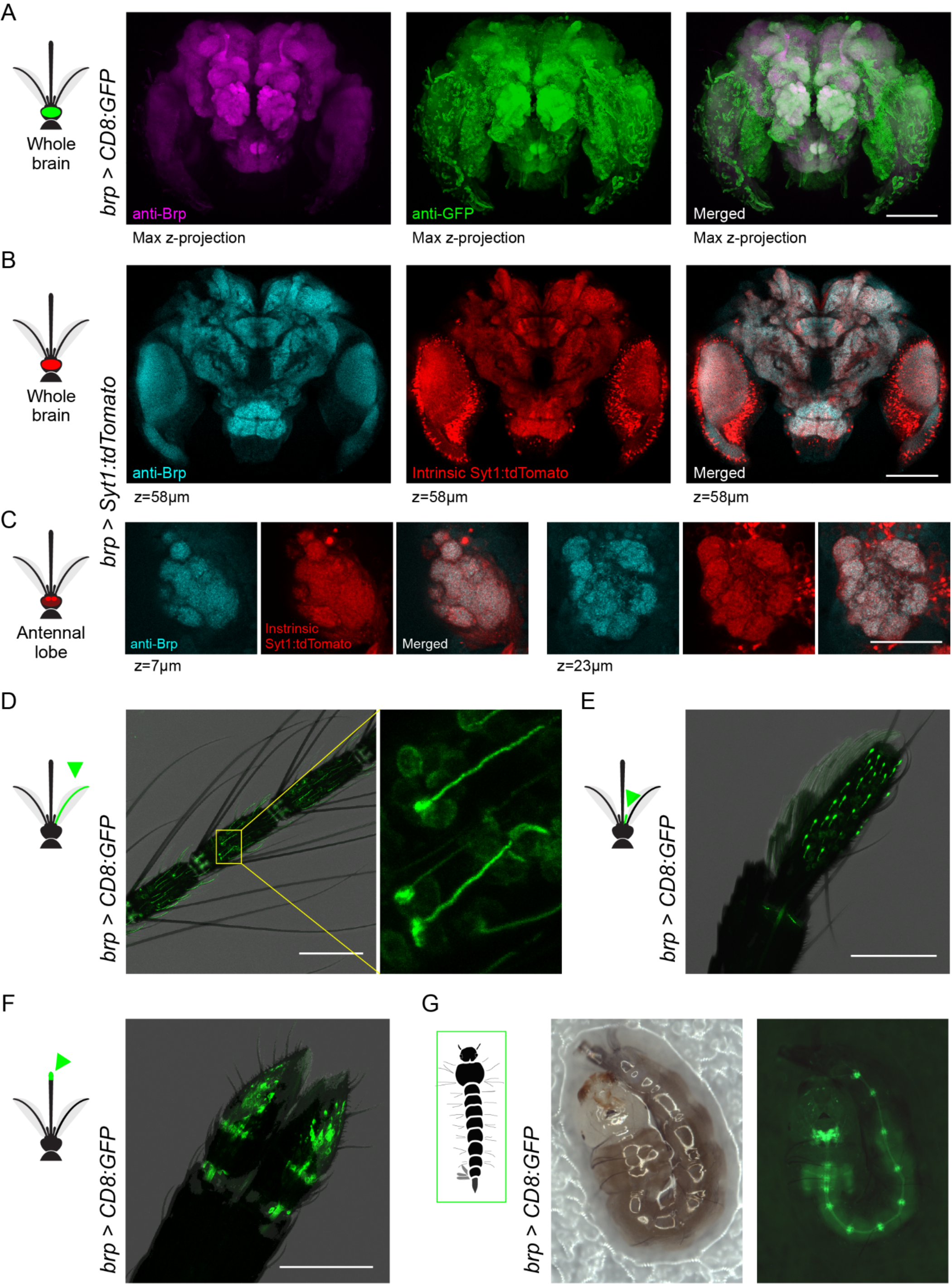
*brp-T2A-QF2w* drives expression in the central and peripheral nervous system of *Ae. aegypti* mosquitoes. (**A**) Anti-Brp (neuropil) and anti-GFP staining in brain of *brp>CD8:GFP* female. (**B**,**C**) Anti-Brp and intrinsic Syt1:tdTomato signal in representative z-planes from the brain (**B**) and antennal lobe (**C**) of *brp>Syt1:tdTomato* adult female. (**D-F**) Intrinsic GFP fluorescence in antenna (**D**), maxillary palp (**E**), and labella **(F)** of *brp>CD8:GFP* adult females. Labelled cells in the inset of (**D**) show elongated dendrites characteristic of antennal trichoid sensilla, while those in maxillary palp show capped dendrites characteristic of capitate peg sensilla. **(G)** Intrinsic GFP fluorescence in *brp>CD8:GFP* larva showing signal in brain, ventral nerve cord, and segmental ganglia. The larva moved slightly between the consecutive bright-field (left) and fluorescent (right) images. Scale bars 100 µm, except 50 µm in (**C**).

We tested the efficacy of *brp-T2A-QF2w* as a pan-neuronal driver by crossing it to a *QUAS-CD8:GFP* effector strain (Matthews et al., 2019) and examining the brain of the resulting *brp>CD8:GFP* progeny (Figure 4A). As expected, anti-GFP staining revealed strong labelling in both neuropil and surrounding cell bodies. Since Brp is a presynaptic protein, a reporter that translocates to presynaptic sites would provide an even better test of whether the *brp-T2A-QF2w* driver recapitulates the pattern of endogenous *brp* expression. We therefore used pBac transposition to generate a *QUAS-Syt1:tdTomato* effector strain in which tdTomato is fused to Syt1 and thus should be localized to presynaptic sites. We then crossed this to our driver to generate *brp>Syt1:tdTomato* animals and assessed co-localization of anti-Brp (nc82) signal and intrinsic tdTomato fluorescence in dissected brains. The two signals were strongly co-localized at the scale of both the whole brain (Figure 4B) and antennal lobe glomeruli (Figure 4C).

We also characterized labelling in the peripheral nervous system of *brp>CD8:GFP* mosquitoes, focusing on well characterized chemosensory organs. We observed strong labeling of neurons with the expected dendritic morphology in the antenna (Figure 4D), maxillary palp (Figure 4E), and labella (Figure 4F) of adult females. A companion paper also observed labeling in sensory neurons of the female stylet, a syringe-like set of mouthparts that pierce the skin to draw blood (Jové et al., 2020). We also confirmed expression in the larval nervous system (Figure 4G).

### *Syt1:GCaMP6s* and *brp-T2A-QF2w* have normal life history but show a modest reduction in behavioral performance

Both the pan-neuronal pre-synaptic GCaMP strain and pan-neuronal driver strain were viable and easy to breed in the lab. However, more subtle effects of these genetic manipulations on fitness should be considered when studying behavior and interpreting neurobiological studies. We therefore quantified several key life history traits in heterozygotes of both strains after 6 or more generations of outcrossing to wildtype Orlando mosquitoes (Materials and methods). Note that we have not attempted to make either strain homozygous and foresee most downstream testing taking place in heterozygotes.

We first examined the rate at which third to fourth instar larvae inherit each transgenic construct from a heterozygous parent and found no significant deviations from the expected 0.5 for Mendelian traits (one-sample t-test, *P*>0.05, Figure 5A). This finding confirms that neither knock-in causes a substantial reduction in embryonic or early larval survival in the laboratory. We next examined sex ratio, larval-adult survival, blood-feeding rate, oviposition rate, and fecundity. In all cases, the pan-neuronal lines were comparable to wildtype controls, although *brp-T2A-QF2w* showed a marginal trend for reduced blood-feeding and oviposition (Figure 5B-F).

**Figure 5.**
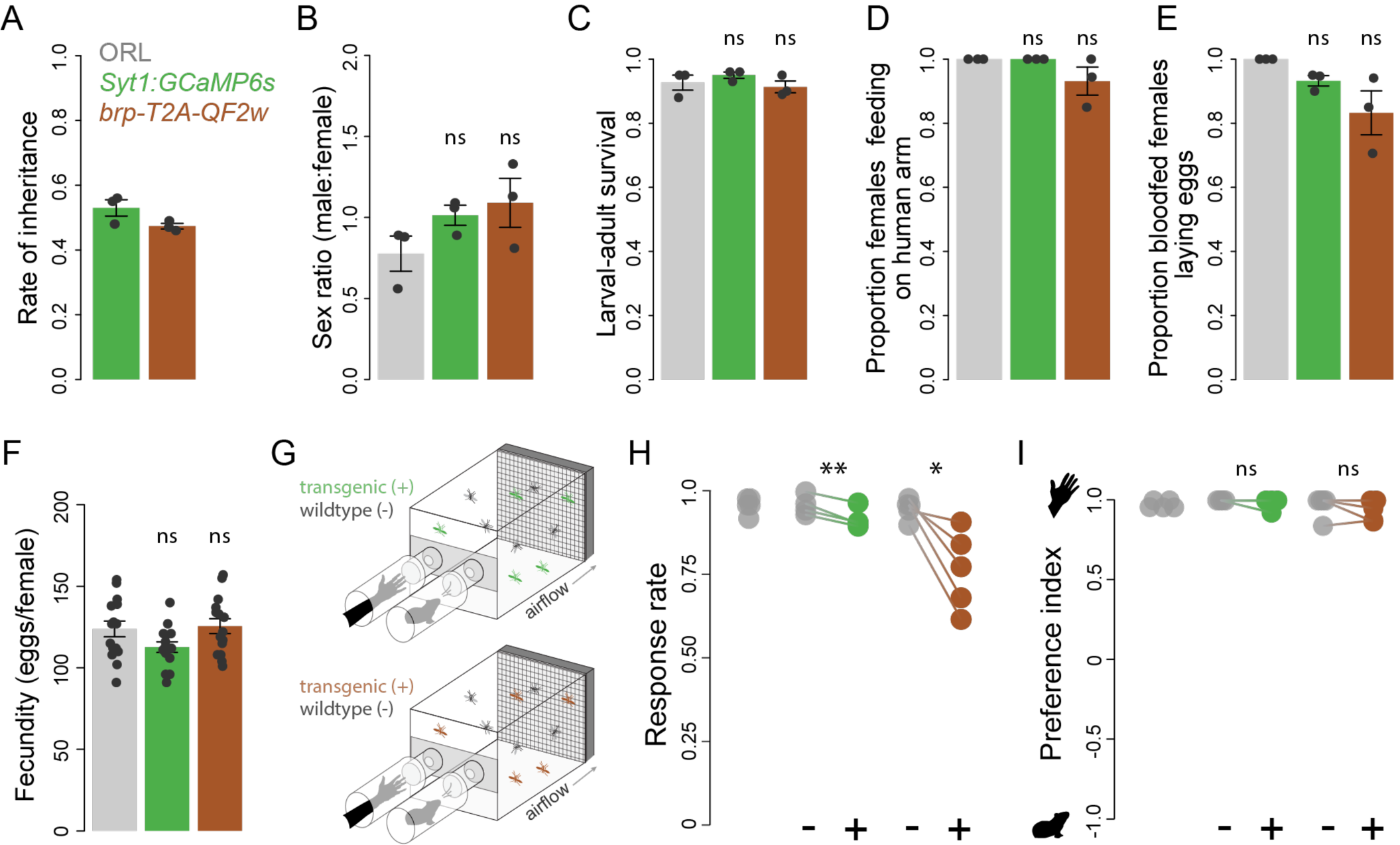
Life history and olfactory behavior of pan-neuronal strains. *Syt1:GCaMP6s* (green) and *brp-T2A-QF2w* (brown, no effector present) heterozygotes were tested alongside wildtype Orlando (grey) mosquitoes. (**A**) Transgenic constructs were inherited by ∼50% of offspring in heterozygote x wildtype crosses (one-sample *t*-test for µ=0.5, *P*>0.05). (**B-F**) Transgenic lines resembled wildtype in key life history traits, including sex ratio (**B**), larval-adult survival (**C**), blood-feeding rate (**D**), oviposition rate (**E**), and fecundity (**F**). Bars and lines indicate mean ± SEM. Dots in (**A-E**) show biological replicates (n=3 groups per strain where each group included the offspring of 5 females). Dots in (**F**) indicate individual females (n=15, including 5 from each of the three groups tested in (**A-E**)). Pairwise *t*-tests comparing each transgenic to wildtype for (**B**,**F**) (raw values) and (**C-E**) (logit transformed values) were all *P*>0.05. (**G**) Schematic of two-port olfactometer assays used to test preference for human *versus* guinea pig odor. Transgenic heterozygotes were tested alongside their wildtype siblings in a paired design. The wildtype ORL strain was also tested separately (not depicted). (**H**,**I**) Response rate (**H**) and preference index (**I**) from preference trials. Lines connect data points for sibling groups from the same trial. Paired t-test: ns, not significant; *, *P*<0.05; **, *P*<0.01.

Finally, we compared the host odor response and preference of heterozygous knock-ins to wildtype. Females were given 6 minutes to choose between human and guinea pig odor in a two-port olfactometer (Figure 5G, Materials and methods). Interestingly, both pan-neuronal lines showed significant reductions in the number of females responding to either stimulus. The reduction was minor for *Syt1:GCaMP6s* (4%), but more substantial for *brp-T2A-QF2w* (19%) (Figure 5H), suggesting that the latter may experience a general decrease in behavioral motivation or performance. Nevertheless, responding females of both genotypes still strongly preferred human odor over the odor of a non-human animal (Figure 5I), indicating normal olfactory discrimination.

## Discussion

We have developed two pan-neuronal genetic tools in *Ae. aegypti* mosquitoes by targeted knock-in of effectors or transcriptional activators to the native locus of broadly expressed neural genes. One strain expresses a Syt1:GCaMP6s fusion protein without amplification and is suitable for synaptic imaging (*Syt1:GCaMP6s*, Figure 2B-C). The other strain is a flexible driver that can be used in concert with the Q binary system to drive strong expression of diverse effector molecules for myriad applications (*brp-T2A-QF2w*, Figure 3C, Figure 4). Both strains are easy to breed in the laboratory and resemble wildtype mosquitoes in an array of life history traits and in an olfactory discrimination task (Figure 5). However, the *brp-T2A-QF2w* driver showed a moderate reduction in overall behavioral motivation/response rates, which should be kept in mind when designing and interpreting experiments.

Pan-neuronal expression is defined as expression in all neurons and no other cell types. Confocal imaging of reporter signal was consistent with this expectation. However, a more sensitive test for pan-neuronal expression of the *brp-T2A-QF2w* driver would be to confirm co-localization with a protein found in the soma of all and only neurons and to confirm lack of co-localization with a protein found in the soma of glia but not neurons. This would allow one to more carefully inspect individual cells within a focal brain region or peripheral tissue and determine whether an effector was missing from any neurons or present in any glia. Unfortunately, we were not able to obtain reliable results with commercially available neuronal or glial antibodies from *Drosophila* (anti-Elav, anti-Repo). Further verification is thus necessary for applications where precise neuron counts are important. Advances in transgenics, *in situ* hybridization protocols, and antibody development will facilitate validation studies in the future.

We tried a variety of genetic approaches while working to generate the pan-neuronal tools presented here and hope that others may learn from both our successes and failures. For one, while promoter fusions rarely capture the pattern of expression of target genes in *Ae. aegypti*, it is now clear that in-frame knock-in of T2A-effector constructs offers a reliable alternative. This strategy has now worked for four sparsely-expressed sensory receptors (*ppk301*, Matthews et al., 2019; *Gr4, Ir7a, Ir7f*, Jové et al., 2020), two broadly expressed synaptic genes (*Syt1* and *brp*, this paper), and at least four other genes (McBride lab unpub. data). The approach may also be useful in other non-model species with large genomes where regulatory elements are distributed across long intergenic regions, as long as CRISPR/Cas9-mediated homology directed repair is working efficiently (Kistler et al., 2015; Li et al., 2017).

When developing pan-neuronal drivers, it is also important to consider how broad expression of exogenous proteins may interfere with endogenous processes and cause toxicity (Rezával et al., 2007; Riabinina et al., 2015; Steinmetz et al., 2017). We found that *Syt1-T2A-QF2-QUAS-GCaMP6s* was larval lethal, likely due to overexpression of QF2 and/or GCaMP6s. This problem was ultimately solved by one or both of two modifications: we used a weaker version of the QF2 transcriptional activator (QF2w; Riabinina et al., 2015) and targeted a less highly expressed neural gene (*brp*; Figure 1B) in order to generate the viable *brp-T2A-QF2w* strain (Figure 3C). We also split the binary system in a more conventional way such that the *brp* driver line alone would not express a GCaMP effector. However, crosses between the *brp* driver and various effectors, including a *QUAS-GCaMP6s* strain, also showed normal development (data not shown). Thus, the identity of the transcriptional activator and the expression level of the target gene are important factors to consider when generating broadly expressed drivers.

Looking forward, we hope that the tools presented here will accelerate neurobiological research in *Ae. aegypti*. Pan-neuronal reagents are a cornerstone of any neurogenetic toolkit, offering a global view of neural processes and an entry point for the dissection of the circuitry underlying behavior. Our *brp-T2A-QF2w* driver line has already been used to characterize taste neurons in the stylet, revealing elegant sensory mechanisms female mosquitoes use to distinguish blood from nectar and initiate blood-feeding behavior (Jové et al., 2020). In pilot experiments, the *Syt1:GCaMP6s* also proved useful for recording odor-evoked responses in olfactory glomeruli of the antennal lobe. We expect the continued application of these tools in mosquito research will provide new insight into the neurobiology of behavior and inform the development of new mosquito control strategies.

## Materials and methods

### Ethics and regulatory information

The use of non-human animals in olfactometer trials was approved and monitored by the Princeton University Institutional Animal Care and Use Committee (protocol #1999-17 for live guinea pigs). The participation of human subjects in olfactometer trials was approved by the Princeton University Institutional Review Board (protocol #8170). All human subjects gave their informed consent to participate in work carried out at Princeton University. Human-blood feeding conducted for mosquito colony maintenance did not meet the definition of human subjects research, as determined by the Princeton University IRB (Non Human-Subjects Research Determination #6870).

### Mosquito rearing and colony maintenance

All mosquitoes used in this research were reared at 26°C, 75% RH on a 14:10 light/dark cycle. Larvae were hatched in deoxygenated water and fed Tetramin Tropical Tablets (Pet Mountain, 16110M). Pupae were transferred to plastic bucket or bugdorm cages and provided access to 10% sucrose solution *ad libitum*. Females were allowed to blood-feed on a human arm prior to egg collection. Eggs were collected on wet filter papers (Whatman, 09-805B) and dried for one week before storage.

### CRISPR/Cas9 transgenesis overview

All CRISPR/Cas9 injection mixes were prepared according to the protocols described below and injected into the Orlando (ORL) strain at the Insect Transformation Facility at the University of Maryland Institute for BioScience & Biotechnology.

#### sgRNA design, synthesis, and testing

We designed sgRNAs to target the last coding exons of *Syt1, brp, nSyb, and elav* (as close as possible to the stop codon, 3-5 sgRNAs per gene) using CHOPCHOP (Montague et al., 2014) and CRISPR Design (http://crispr.mit.edu) and prepared them as previously described (Kistler et al., 2015). Briefly, we generated double-stranded DNA template for transcription of each sgRNA via a template-free polymerase chain reaction (PCR) with partially overlapping primers (IDT, PAGE purified) and purified the product with Ampure XP beads (Beckman Coulter, A63880). We then transcribed sgRNAs *in vitro* using the MEGAscript T7 Transcription Kit (ThermoFisher Scientific, AM1334) with 37°C incubation for 12-16 hours, treated them with DNAse at 37°C for 15 min, and purified with MEGAclear Transcription Clean-Up Kit (ThermoFisher Scientific, AM1908). We mixed 3-5 sgRNAs, each targeting a different gene (80 ng/uL each) with Cas9 protein (300 ng/µL; PNA Bio, CP01-200) and injected into ORL embryos. We extracted DNA from groups of injected G0s (n=3∼5 pupae per group, n=3 groups per injection) and amplified the region surrounding each cut site via PCR. We then inferred cutting efficiency by quantifying the rate of small insertion and deletion mutations in the amplified region via fragment length analysis (Carrington et al., 2015) or Illumina MiSeq sequencing (Kistler et al., 2015). For fragment analysis, PCR primers were conjugated with a fluorescent dye and amplicons were submitted to GENEWIZ. For sequencing, the PCR primers contained MiSeq index/adapter sequences and amplicons were sequenced to a depth of ∼2500X.

#### Knock-in injection mix protocol 1

We prepared injection mixes for *Syt1-T2A-QF2-QUAS-GCaMP6s, Syt1-T2A-3XGCaMP6s* according to Kistler et al., 2015. sgRNAs were generated as described above. We prepared donor plasmids using the InFusion HD Kit (Clontech, 638910) and EndoFree Plasmid Maxi Kit (Qiagen, 12362), and verified them via Sanger sequencing. We mixed donor plasmid (700 ng/uL) and sgRNA (80 ng/uL), purified the mixture via ethanol precipitation, and resuspended in Ambion nuclease-free water (Life Technologies, AM9937), before adding recombinant Cas9 protein (300 ng/µL; PNA Bio, CP01-200) for embryo injection.

#### Injection mix protocol 2

We prepared injection mixes for *Syt1-T2A-GAL4d-UAS-GCaMP6s, Syt1:GCaMP6s*, and *brp-T2A-QF2w* according to Matthews et al., 2019. Briefly, we generated DNA template for transcription of a new batch sgRNA (separate from that used for validation) by annealing two partially overlapping PCR primers (IDT, PAGE purified) and extending with NEBNext High-Fidelity polymerase (NEB, M0541S). We then transcribed each sgRNA *in vitro* using the HiScribe T7 Kit (NEB, E2040S) with 37°C incubation for 6 hours before treating with DNase at 37°C for 15 min, purifying with RNAse-free SPRI beads (Agencourt RNAclean XP, Beckman-Coulter A63987), and eluting in Ambion nuclease-free water (Life Technologies, AM9937). We prepared donor plasmids using the InFusion HD Kit (Clontech, 638910) and NucleoBond Xtra Midi EF Kit (Macherey-Nagel, 740420.10) and verified them via Sanger sequencing. We directly mixed recombinant Cas9 protein (300 ng/µL; PNA Bio, CP01-200), sgRNA (80 ng/µL) and donor plasmid (700 ng/uL) for embryo injection.

#### Breeding and screening

We hatched G0 embryos in diluted hatching broth 3 days post-injection, and then collected hatchlings and replaced the hatching broth every day thereafter for approximately 2 weeks. At the pupal stage, we separated female and male G0s, and crossed them to wildtype ORL *en masse* (for female G0s) or in single pairs or small groups (for male G0s, to maximize the number that had an opportunity to mate). Mated ORL and G0 females were blood-fed and transferred to oviposition vials to lay eggs (n=1-6 females per vial). We hatched G1 egg papers separately in small larval pans, so all positive individuals in a pan were most likely from the same founder. We screened larvae 4-5 days post-hatching for 3XP3-dsRed phenotype (red fluorescence in the eyes and optic nerves) under an epifluorescence scope. Positive individuals were reared and outcrossed to wildtype ORL for further breeding and validation. The only exception to these procedures was for *Syt1-T2A-QF2-QUAS-GCaMP6s* and *Syt1-T2A-3XGCaMP6s*, where mating and egg collection for the cross between male G0s and wildtype ORL females occurred in large cages *en masse*, and G1 progeny were hatched and screened in larger groups without knowledge of family membership. We validated insertions by Sanger sequencing of PCR amplicons that stretched from within the donor construct to outside of one homology arm. We outcrossed validated lines to ORL for at least 6 generations to minimize potential effects of off-target cutting by Cas9.

### Syt1 targeting overview

We used the same sgRNA and donor plasmid homology arms (left: 1374 bp, right: 1821 bp) for all *Syt1* knock-in injections. The sgRNA (GCACACTCTTAAAGACCCGG<u>AGG</u>, PAM site underlined) targeted a cut site 21 bp upstream of the stop codon. To preserve the *Syt1* coding sequence, the final seven codons of *Syt1* downstream of the cut site were included in the donor plasmid 5’ of the T2A motif, with synonymous codon substitutions incorporated to protect the sequence from Cas9 cleavage and minimize homology between the plasmid insert and the targeted locus.

### Syt1-T2A-3XGCaMP6s screening and donor plasmid details

We injected 1098 ORL embryos and obtained two 3XP3-dsRed-positive G1 families among the offspring of G0 females. However, neither family was positive at G2, probably due to lack of genomic integration of the donor construct. We then screened offspring from G0 males (crossed and oviposited *en masse*) and found 51 positive G1 larvae. We selected two larvae for further outcrossing and validation. PCR verification revealed that one had integrated the full insert with three copies of GCaMP6s, while the other had an insert with only two copies, possibly due to unexpected recombination events between the tandem GCaMP6s sequences. We proceeded to characterize expression for the 3X insert only.

#### Syt1LeftArm-T2A-GCaMP6s-T2A-GCaMP6s-T2A-GCaMP6s-SV40-3XP3-dsRed-SV40-Syt1RightArm

[1] Plasmid backbone from psL1180, linearized with restriction enzymes NsiI-HF (New England Biolabs #R3127S) and AvrII (New England Biolabs #R0174S).

[2] *Syt1* left homology arm from ORL genomic DNA (NCBI LOC5565901) with final seven *Syt1* codons included in the reverse primer (underlined) (Primers: Forward, 5’-caggcggccgccataGCCGGC CTTCTGATAACTGATACCAG-3’,lowercase indicates the homology sequence needed for InFusion cloning; Reverse, 5’-ccctctcccgatcc<u>ATC TTTCTTATCATCTTCTGG</u>GTCTTTAAGAGTGT GCCATTGTGCG -3’).

[3] *Syt1* right homology arm from ORL genomic DNA (NCBI LOC5565901) (Primers: Forward, 5’-tagcggtcgtcctagCGGAGGACGACAAGAAGGACT AAGG-3’; Reverse, 5’-tattaataggcctagCCTCTAC TTTCCAATATATTGTCCGGAATCG-3’).

[4] T2A-GCaMP6s from pGP-CMV-GCaMP6s (Addgene plasmid #40753) with T2A sequence includedintheforwardprimer(underlined) (Primers: Forward, 5’-caggcggccgccataTGCATG GATCGGGA<u>GAGGGCCGCGGCTCCCTGCTGA CCTGCGGCGACGTGGAGGAGAACCCCGGCC CC</u>ATGGGTTCTCATCATCATCATCATCATGG-3’; Reverse, 5’-TCACTTCGCTGTCATCATTTGTACAAACTC -3’).

[5] SV40-3XdsRed-SV40 from psL1180-Or4-GSG-T2A-mCD8GFP-3XP3dsRed (McBride Lab) (Primers: Forward, 5’-atgacagcgaagtgaCAAATAA CGGCCGCGACTCTAG-3’; Reverse, 5’-ttaataggc ctaggaCGACCGCTAAGATACATTGATGAG-3’).

### Syt1-T2A-QF2-QUAS-GCaMP6s screening and donor plasmid details

We injected 1200 ORL embryos and obtained five 3XP3-dsRed-positive G1 families among the offspring of G0 females. All larvae from two families had strong green fluorescence in the central and peripheral nervous system, but died at the late larval stage. The other three families were not positive at G2.

#### Syt1LeftArm-T2A-QF2-Hsp70-5XQUAS-GCaMP6s-SV40-3XP3-dsRed-SV40-Syt1RightArm

[1] Plasmid backbone from *psL1180, linearized* with restriction enzyme NsiI-HF (New England Biolabs #R3127S).

[2] QF2-Hsp70 from pattB-synaptobrevin-7-QFBDAD-hsp70 (Addgene plasmid #46115) (Primers: Forward, 5’-ATGCCACCCAAGCGCAA AACG-3’; Reverse, 5’-ccaagcttggatccaGGCCGC GGATCTAAACGAGTTT-3’).

[3] 5XQUAS from pQUAST-mCD8-GFP (Addgene plasmid #24351) (Primers: Forward, 5’-caggcggc cgccataTGCATGGATCCAAGCTTGGATCCGGG T-3’; Reverse, 5’-atgatgagaacccatTTTGCTCGA GCCGCGGCC-3’).

[4] Syt1LeftArm-T2A and GCaMP6s-SV40-3XP3-dsRed-SV40-Syt1RightArmfromSyt1LeftArm-T2A-GCaMP6s-T2A-GCaMP6s-T2A-GCaMP6s-SV40-3XP3-dsRed-SV40-Syt1RightArm (above).

### Syt1-T2A-GAL4d-UAS-GCaMP6s screening and donor plasmid details

We injected 1500 ORL embryos and obtained four 3XP3-dsRed-positive G1 families among the offspring of G0 females. Two families were still positive at G2, but we were not able to validate integration via PCR. We therefore injected another 2000 ORL embryos and obtained four positive G1 families among the offspring of G0 females. Two families were again positive at G2 and PCR followed by Sanger sequencing revealed proper genomic integration for one family.

#### Syt1LeftArm-T2A-GAL4d-Hsp70-3XUAS-GCaMP6s-SV40-3XP3-dsRed-SV40-Syt1RightArm

[1] Plasmid backbone from psL1180, linearized with restriction enzymes NsiI-HF (New England Biolabs #R3127S) and AvrII (New England Biolabs #R0174S).

[2] GAL4d from pBPGAL4.1Uw (Addgene plasmid #26226)(Primers:AD domain,forward,5’-caggcggccgccataTGCATGGATCCGCCAACTTCAACCAGAGTGG-3’, reverse, 5’-tatcgatagacgtca CTACTCCTTCTTTGGGTTCGG-3’; BD domain, forward,5’-ATGAAGCTGCTGAGTAGTATTGA AC-3’, reverse 5’-aagttggcggatccaGATACCGTC AGTTGCCGTTGAC-3’).

[3] 3XUAS-GCaMP6s-SV40 from *Syt1LeftArm-T2A-GCaMP6s-T2A-GCaMP6s-T2A-GCaMP6s-SV40-3XP3-dsRed-SV40-Syt1RightArm*(above) with 3XUAS included in the forward primer (underlined) (Primers: Forward, 5’-caggcggccgcc ataTGCAT<u>CGGAGTACTGTCCTCCG</u>ag<u>CGGAG TACTGTCCTCCG</u>ag<u>CGGAGTACTGTCCTCCG</u>A AGCTTGATATCGAATTCCTGCAG-3’;Reverse, 5’-TCACTTCGCTGTCATCATTTGTACAAACTC - 3’).

[4] Syt1LeftArm-T2A, Hsp70 and 3XP3-dsRed-SV40-Syt1RightArm from Syt1LeftArm-T2A-QF2-Hsp70-5XQUAS-GCaMP6s-SV40-3XP3-dsRed-SV40-Syt1RightArm (above).

### Syt1:GCaMP6s screening and donor plasmid details

We used the 3XGS linker (GGATCGGGCTCCG GCTCC) to produce a fused Syt1:GCaMP6s protein that should be translocated and concentrated in presynaptic sites. We injected 1200 ORL embryos and obtained two 3XP3-dsRed-positive G1 families among the offspring of G0 females. Both families were positive at G2, one of which was validated through PCR and Sanger sequencing. Donor plasmid is available at Addgene (#159636).

#### Syt1LeftArm-3XGS-GCaMP6s-SV40-3XP3-dsRed-SV40-Syt1RightArm

[1] Plasmid backbone from *Syt1LeftArm-T2A-GAL4d-Hsp70-3XUAS-GCaMP6s-SV40-3XP3-dsRed-SV40-Syt1RightArm*(above),linearized with inverse PCR to remove the T2A-GAL4d-Hsp70-3XUASelements.Oneprimeralso included the 3XGS linker (underlined). (Primers: Forward, 5’-ATGGGTTCTCATCATCATCATCAT CATGG-3’; Reverse, 5’-<u>GGAGCCGGAGCCCGA TCC</u>ATCTTTCTTATCATCTTC-3’).

[2] Self-ligation with T4 ligase to generate the donor plasmid.

### brp-T2A-QF2w overview and details

We used a sgRNA (GCAACTGGTACAGATGAC AC<u>AGG</u>, PAM site underlined) that targeted a cut site in the penultimate exon of the *brp* gene, 46 codons upstream from the stop codon, which resides in the final exon. To preserve the *brp* coding sequence, the final 46 codons of *brp* downstream of the cut site were therefore included in the donor plasmid 5’ of the T2A motif. This 46 codon fragment was synthesized with synonymous codon substitutions to protect the sequence from Cas9 cleavage and to minimize homology between the plasmid insert and the targeted locus (IDT, gBlocks, sequence below). Donor plasmid is available at Addgene (#141094).

We injected 1533 ORL embryos and obtained six 3XP3-dsRed-positive G1 families among the offspring of G0 females. Two of these families were positive at G2, and one was validated via PCR and Sanger sequencing.

#### brpLeftArm-T2A-QF2w-Hsp70-3XP3-dsRed-SV40-brpRightArm

[1] Plasmid backbone from psL1180, linearized with restriction enzymes NsiI-HF (New England Biolabs #R3127S) and AvrII (New England Biolabs #R0174S).

[2] *brp* left homology arm (NCBI LOC5570381) (Primers: Forward, 5’-caggcggccgccataATGAC CGGCTACCATGACCACTTTATAGTA-3’; Reverse,5’-TCATCTGTACCAGTTGCAGTAAACGTTCC-3’).

[3] *brp* right homology arm (NCBI LOC5570381) (Primers: Forward, 5’-tgtatcttatcctagCACAGGAA GAGCAGAACCAAAAAGAAAAGAC-3’; Reverse, 5’-tattaataggcctagTTTCGAATCTGTGACAAATTTCCCGATAAGAACT-3’).

[4] QF2w-Hsp70from pQF2wWB(Addgene plasmid #61313) (Primers: Forward, 5’-GCAAAA CGCTTAACGCTGCG-3’, Reverse, 5’-cgtaggata acttcgGGATCTAAACGAGTTTTTAAGCAAACT-3’)

[5] *brp* synthetic fragment with synonymous codon substitutions: AACTGGTACAGATGACCCAGGAAGAACAGAACCAGAAGGAAAAGACCATCATGGATCTGCAGCAGGCCCTGAAGAACGCCCAGGCCAAGCTGAAGACCGCCCAGTCGCAGCCGCAGGATGCCGGACCGGCCGGATTCCTGAAGTCGTTCTTTGGATCGGGAGAGGG

[6] Sequences of T2A and SV40-dsRed-SV40 same as in Syt1 constructs above.

### Generation of *QUAS-Syt1:tdTomato* via pBac-mediated transposition

This line was generated in the ORL background with pBac-mediated transposition according to a previously published method that used pBac mRNA (Jové et al., 2020). Briefly, we generated the template for *in vitro* transcription of pBac mRNA via PCR amplification from a plasmid containing the pBac coding sequence (a gift from Leslie Vosshall; primers forward 5’-GAAACTAA TACGACTCACTATAGGGAGAGCCGCCACATGGGTAGTTCTTTAGACGATG-3’, reverse 5’-CTTA TTAGTCAGTCAGAAACAAC-3’).ThePCR amplicon was purified using RNAse-free SPRI beads (Agencourt RNAclean XP, Beckman-Coulter A63987) and then used for *in vitro transcription* with the HiScribe T7 ARCA mRNA Kit (with tailing, NEB, E2060S). Transcription products were purified using RNAse-free SPRI beads, and eluted in Ambion nuclease-free water (Life Technologies, AM9937). The transgene plasmid was generated using the InFusion HD Kit (Clontech, 638910) and the NucleoBond Xtra Midi EF Kit (Macherey-Nagel, 740420.10). A mixture of plasmid (500 ng/uL) and pBac mRNA (300 ng/uL) was injected into 346 ORL embryos. We obtained 14 3XP3-ECFP-positive families among G1 larvae. Eleven families remained positive at G2. We outcrossed positive families to ORL for 3 generations and then randomly chose one family to cross with the *brp-T2A-QF2w* driver line.

#### pBacLeftArm-15XQUAS-Syt1:tdTomato-SV40-3XP3-ECFP-SV40-pBacRightArm

[1] Plasmid backbone from *pBacLeft-15XQUAS-Syt1:GCaMP6s-SV40-3XP3-ECFP-SV40-pBacRight* (a gift from Leslie Vosshall), linearized with inverse PCR to remove the GCaMp6s element (Primers: Forward, 5’-GATCTTTGTGAA GGAACCTTACTTCTGTGGTG-3’; Reverse, 5’-CGATCCGGAACCCGATCCGTCTTTCTT -3’).

[2] tdTomato from transgenic *Drosophila melanogaster* that contained tdTomato insertion (a gift from Mala Murthy) (Primers: Forward, 5’-tcgggttccggatcgATGGTGAGCAAGGGCGAGG -3’; Reverse, 5’-tccttcacaaagatcCTTGTACAGCTCGTCCATGCC-3’).

### Characterization of reporter expression

#### Brain immunostaining

Brain immunostaining was carried out as previously described (Matthews et al., 2019). Heads of 7–10 day old mated mosquitoes were fixed in 4% paraformaldehyde (Electron Microscopy Sciences, 15713-S) for 3 hours at 4°C. Brains were dissected in PBS and blocked in normal goat serum (2%, Fisher Scientific, 005-000-121) for 2 days at 4°C. We then incubated brains in primary antibody solution for 2–3 days, followed by secondary antibody solution for another 2–3 days at 4°C. Brains were mounted in Vectashield (Vector, H-1000) with the anterior side facing the objective. Confocal stacks were taken with a 20X lens with XY resolution of 1024×1024 and Z-step size of 1 µm. Primary antibodies: rabbit anti-GFP (1:10,000 dilution, ThermoFisher, A-11122) and mouse NC82 (1:50 dilution, DHSB, AB_2314866). Secondary antibodies: goat-anti-rabbit Alexa 488 (1:500 dilution, ThermoFisher, A27034SAMPLE), goat-anti-mouse CF680 (1:500 dilution, Biotium, 20065-1) and goat-anti-mouse Cy3 (1:500 dilution, Jackson ImmunoResearch, 115-165-062).

#### Brain raw fluorescence

Brains of 7-10 day old mosquitoes were fixed in 4% paraformaldehyde for 30 mins at 4°C and dissected in PBS, before mounting directly in Vectashield for confocal imaging.

#### Peripheral organs

We removed the antenna, maxillary palp, or proboscis of 7–10 day old female mosquitoes with sharp forceps, dipped them in pure ethanol for ∼15 sec, and mounted them on slides in pure glycerol for direct confocal imaging.

#### Larvae

Larvae 4-5 days post-hatching were placed on wet filter paper with the ventral side facing the objective and imaged with an epifluorescence scope.

### Fitness tests

We characterized the fitness and behavior of *Syt1:GCaMP6s* and *brp-T2A-QF2w* mosquitoes alongside wildtype ORL controls. We collected freshly dried eggs from individual heterozygous females crossed to ORL males. For each replicate (n=3 replicates per line), we hatched eggs from five females in a single pan of hatching broth. We then screened 4-day-old larvae for 3XP3-dsRed under an epifluorescence scope and calculated the rate of inheritance (proportion of positive larvae). Survival deficits in embryos and young larvae should result in rates lower than 0.5.

We continued to rear 3XP3-dsRed+ larvae, separated male and female pupae for eclosion, and recorded sex ratio and larva-to-adult survival rate. We recorded the same variables in ORL wildtype families that had been hatched and mock screened under an epifluorescence scope as described above. We then crossed virgin females from each replicate (both knock-ins as well as wildtype) to ORL wildtype males and assessed blood-feeding rates by inserting a human arm (25-year-old East Asian male) into the cage and recording the proportion of females fully engorged after 10 mins. Three days post blood-feeding, we transferred ∼20 individual blood-fed females per replicate into oviplates (Ioshino et al., 2018) and recorded the proportion of females that laid any eggs (oviposition rate). Five egg papers from each replicate were subsequently imaged under a dissection microscope to count the number of eggs per female (fecundity). We only counted eggs for females that laid, making fecundity and oviposition rate estimates independent.

### Olfactometer host-preference assay

We used a two-port olfactometer to test the host preference of female mosquitoes as previously described (McBride et al., 2014). One host port contained a human hand and arm up to the elbow (25-year-old East Asian male), while the other port contained a guinea pig (*Cavia porcellus*; one of two 4–5 year old pigmented females). We used a paired experimental design to directly compare the behavior of knock-in heterozygotes and their wildtype siblings (Figure 4G). For each knock-in line (*Syt1:GCaMP6s* and *brp-T2A-QF2w*), we crossed heterozygotes with ORL wildtype mosquitoes and reared the progeny in a single pan. We expected ∼50% of progeny to carry the knock-in, but did not screen and separate positive and negative siblings. Instead, we reared, housed, and tested siblings in a mixed group. At 6-9 days post-eclosion, we sorted females and housed them in plastic cups overnight with access to water only (no sucrose). Before a given trial, we acclimated 70-100 females (n=70 for *Syt1:GCaMP6s*, n=100 for *brp-T2A-QF2w*) in the olfactometer for 5 min. We then opened a sliding door and activated a fan to pull air through the two host chambers and expose mosquitoes to host odour. Mosquitoes were able to fly upwind, sample the host-odour streams, and enter either host port. After 6 min, we collected responding mosquitoes trapped in each host port and non-responding mosquitoes in the releasing chamber. We froze the mosquitoes at -20°C and screened them for the 3XP3-dsRed phenotype under an epifluorescence scope on the same day. The response rate (#responding / #total) and preference index [(#human - #guinea pig) / #responding] were calculated separately for the 3XP3-dsRed positive individuals and their wildtype siblings. We reared and tested wildtype ORL mosquitoes at the same time and in the same way except females were tested in groups of 50 and did not need to be screened afterwards.

### Reagents and data availability

Donor plasmids for *Syt1:GCaMP6s* and *brp-T2A-QF2w* are available at Addgene (#159636 and #141094). Donor plasmids for other constructs, eggs for *Syt1:GCaMP6s, brp-T2A-QF2w* and *QUAS-Syt1:tdTomato*, and confocal images are available upon request.

## Acknowledgements

We thank Veronica Jové, Leslie Vosshall, and Lu Yang for discussion and comments on the manuscript; Chris Potter for advice on using the Q system; Benjamin Matthews, Meg Younger, Zachary Gilbert, Matthew DeGennaro and members of Aedes Toolkit Group for advice on transgenesis; Rob Harrell for mosquito embryo injections; and Azwad Iqbal for help with mosquito breeding. This work was funded in part by the National Institutes of Health via grants from NIDCD (R00DC012069) and NIAID (DP2AI144246) to C.S.M. C.S.M.’s laboratory is also supported by a Pew Scholar Award, a Searle Scholar Award, a Klingenstein-Simons Fellowship, and a Rosalind Franklin New Investigator Award. C.S.M. is a New York Stem Cell Foundation – Robertson Investigator.

## Author contributions

Z.Z. and C.S.M conceived the project and design experiments. Z.Z. and D.T. generated and tested transgenic lines. All authors helped interpret experiments. Z.Z. and C.S.M. wrote the paper.

